# Hypoxic response is driven by the BAF form of SWI/SNF

**DOI:** 10.1101/2022.02.16.480689

**Authors:** Kathleen Diep Tran, Tomali Chakravarty, Jada Lauren Garzon, Anita Saraf, Laurence Florens, Michael P. Washburn, Arnob Dutta

## Abstract

SWI/SNF has been shown to have important functions in hypoxia-mediated gene expression through roles of its catalytic and core subunits. Since SWI/SNF exists as three distinct assemblies, and usage of complex specific subunits of the complex can be expected to vary within a given cell under changing environmental conditions. It remains an open question as to the compositional makeup of SWI/SNF and the roles of individual complexes in gene expression and cell viability in a hypoxic environment. In our current study, we find that hypoxia regulates levels of unique subunits that define each complex. Protein levels of ARID2 and PBRM1, members of PBAF and BRD9, a member of ncBAF, are downregulated in hypoxic cells, while members of BAF complex are retained. Our studies further show that loss of ARID1A, ARID1B and DPF2, which are unique subunits of BAF, reduces induction of HIF target genes and ARID1A or DPF2 are important for cell survival during hypoxia. Collectively, our results provide evidence that levels of SWI/SNF forms are not static within cells, but can be dynamically altered as a response to environmental changes.

## Introduction

Hypoxia, or low oxygen tension, is a hallmark of many pathological symptoms including ischemia, chronic inflammatory disease, rheumatoid arthritis and cancers. Specifically, in solid tumors, increased oxygen demand during proliferation and growth of tumor mass causes irregular vascularization^1,2^. This leads to improper oxygen distribution, creating a milieu of hypoxic tumor cells^3^. Hypoxia in tumor microenvironments induces cellular changes that make these cells highly resistant to chemotherapy, immune and radiation therapy^4,5^. Hypoxia, although associated with conditions such as tumorigenesis and disease, also occurs in normal activity in human physiology and development, including presence in stem cell niches^6^, and being required for the maintenance of hematopoietic^7^, bone marrow, adipocyte^8^ and brain stem cells^9^ in their undifferentiated states. For survival during hypoxia, cells induce several changes impacting regulation of gene expression, translation, and metabolic processes: most prominently a shift from oxidative phosphorylation to glycolysis^10^. This is often achieved by a complex cell signaling network including but not limited to, the HIF^11^, PI3K^12^, MAPK^13^, and NFĸB^14^ pathways, which interact with each other for cell survival. Hypoxia-inducible factor 1 (HIF-1) is a heterodimeric transcription factor that consists of HIF-1β which is constitutively expressed and HIFα (HIF-1α and HIF-2α), the oxygen-regulated subunits. HIFα is normally marked for proteasomal degradation by the von Hippel-Lindau (VHL) complex in oxygen rich environments^15^. However, when oxygen levels begin to diminish, HIFα proteins are stabilized and can induce transcription of genes with adaptive functions^16^. HIF-1α and HIF-2α binding patterns are shown to span around promoters and enhancers of HIF target genes in which the binding sites are dependent on histone modifications and transcription factors^17^. Several transcription factors are needed for the induction of hypoxic genes including CBP and p300^18^, PKM2^19^ and SWI/SNF^11^. Given the important role of hypoxia in both normal physiology and in disease, understanding gene regulatory pathways and usage of chromatin remodelers like SWI/SNF for altered transcription during hypoxia is one of significant importance.

The ATP-dependent chromatin remodeler, SWI/SNF, is highly conserved throughout the eukaryotic kingdom, where this protein complex functions to regulate gene expression levels by modulating chromatin structure^20–24^. The composition of mammalian SWI/SNF contains up to 15 subunits which can be assembled from approximately 29 genes, allowing for variations of the complex across tissues and within a given cell^25^. In mammals, there exist three distinct assemblies: BRG1/BRM-associated factor complexes (BAF), polybromo-associated BAF complexes (PBAF), and non-canonical BAFs (ncBAF)^26,27^. These assemblies share a set of core components, including BAF155/BAF170, BAF60a, BAF57, and INI1 (found in BAF and PBAF only) and catalytic ATPases BRG1 or BRM. Each complex also contains several exclusive subunitscthat distinguish the assemblies, including ARID1A/ARID1B and DPF2 of BAF, ARID2, PBRM1, BRD7 and PHF10 of PBAF, and GLTSCR1/L and BRD9 of ncBAF. The heterogeneity of SWI/SNF allows for its specialized roles in tissue and developmental-specific gene regulation. Genomic studies have shown that the complexes have overlapping roles as BAF has been found mostly at enhancers^28,29^, PBAF can be found at promoter sequences^30,31^, and ncBAF has shown to be found at both enhancers and promoters^32,33^. However, each complex also has their distinct functions, generated by exclusive subunits in each complex. For example, during neural development, PHF10 and ACTL6A subunits will switch to DPF1/3 and ACTL6B, respectively, to allow for differentiation^34,35^. Also, within one complex, paralogs can have opposing roles, like that of ARID1A and ARID1B of BAF in cell-cycle regulation^36^. Further, alterations or inactivation of genes encoding subunits of SWI/SNF have been implicated in over 20% of cancers, causing dysregulation in gene expression^37,38^. These mutations have been shown to be drivers of tumor formation in which there is insufficient oxygen available, creating a hypoxic environment. Studies have shown important functions for SWI/SNF in the hypoxic response. This includes the catalytic subunits, BRG1 and BRM, and accessory components, BAF155/BAF170 and BAF57^39^. Since these subunits are common to the various forms of SWI/SNF, the contribution of each of the distinct forms of SWI/SNF to hypoxic gene expression is not well understood.

In the present study, we have investigated the composition and stoichiometry of the various forms of SWI/SNF and their roles in gene regulation and cell survival as a response to hypoxia. We find that cells maintain high levels of the BAF form of SWI/SNF, with a significant down regulation of subunits of the PBAF complex, along with moderate loss of members of ncBAF. Our studies also show that the alteration in levels of the various forms of SWI/SNF within hypoxic cells coincides with dependence on BAF subunits ARID1A, ARID1B, and/or DPF2 for gene expression and cell survival as a response to hypoxia. Our studies cumulatively point to a fundamental reorganization of stoichiometry of the various forms of SWI/SNF complexes for cellular hypoxic response with primary dependence on the BAF form of SWI/SNF.

## Results

### Hypoxia alters levels of complex specific subunits of SWI/SNF

Studies have shown that SWI/SNF catalytic subunits BRG1/BRM and core subunit members BAF155/BAF170 and BAF57 are important for hypoxic gene expression, but the compositional diversity of SWI/SNF within hypoxic cells has not been well defined. To investigate if hypoxia affected protein levels of members of distinct SWI/SNF complexes, we first examined levels of SWI/SNF subunit proteins by western blotting in nuclear extracts of HCT116 (colorectal carcinoma) cells grown with or without hypoxia (1% O_2_) for 24 hours. We verified that hypoxia was induced by detection of HIF-1α (Figure 1A). Compared to normoxia, the levels of catalytic subunit BRG1 and core components, BAF155, BAF60a, INI1, and BAF57, remained unchanged in hypoxia (Figure 1A). Levels of proteins of unique subunits of BAF, specifically ARID1A, ARID1B, and DPF2, also remained unchanged between the two conditions. However, when we analyzed unique subunits of PBAF, ARID2 and PBRM1, and of ncBAF, BRD9, hypoxic exposure significantly reduced their protein levels (Figure 1A). To confirm our observations, we quantified the intensity of the bands from three independent experiments (Figure 1B). To further confirm that these observations were not cell line specific but were a general response to hypoxia, we analyzed the levels of SWI/SNF subunits in ES-2 (ovarian carcinoma) cells. ES-2 cells displayed a similar alteration of SWI/SNF complex stoichiometry during hypoxia as HCT116, with a decrease in expression levels of ARID2, PBRM1, and BRD9 in hypoxia and maintenance of ARID1A, ARID1B and DPF2, along with catalytic and core members of SWI/SNF (Supplementary Fig. 1A, B). These results suggest that as a response to hypoxia, subunits of PBAF and ncBAF are downregulated while BAF subunits remain unchanged, indicating that BAF subunits may play an important role in hypoxic gene expression.

**Figure 1.**
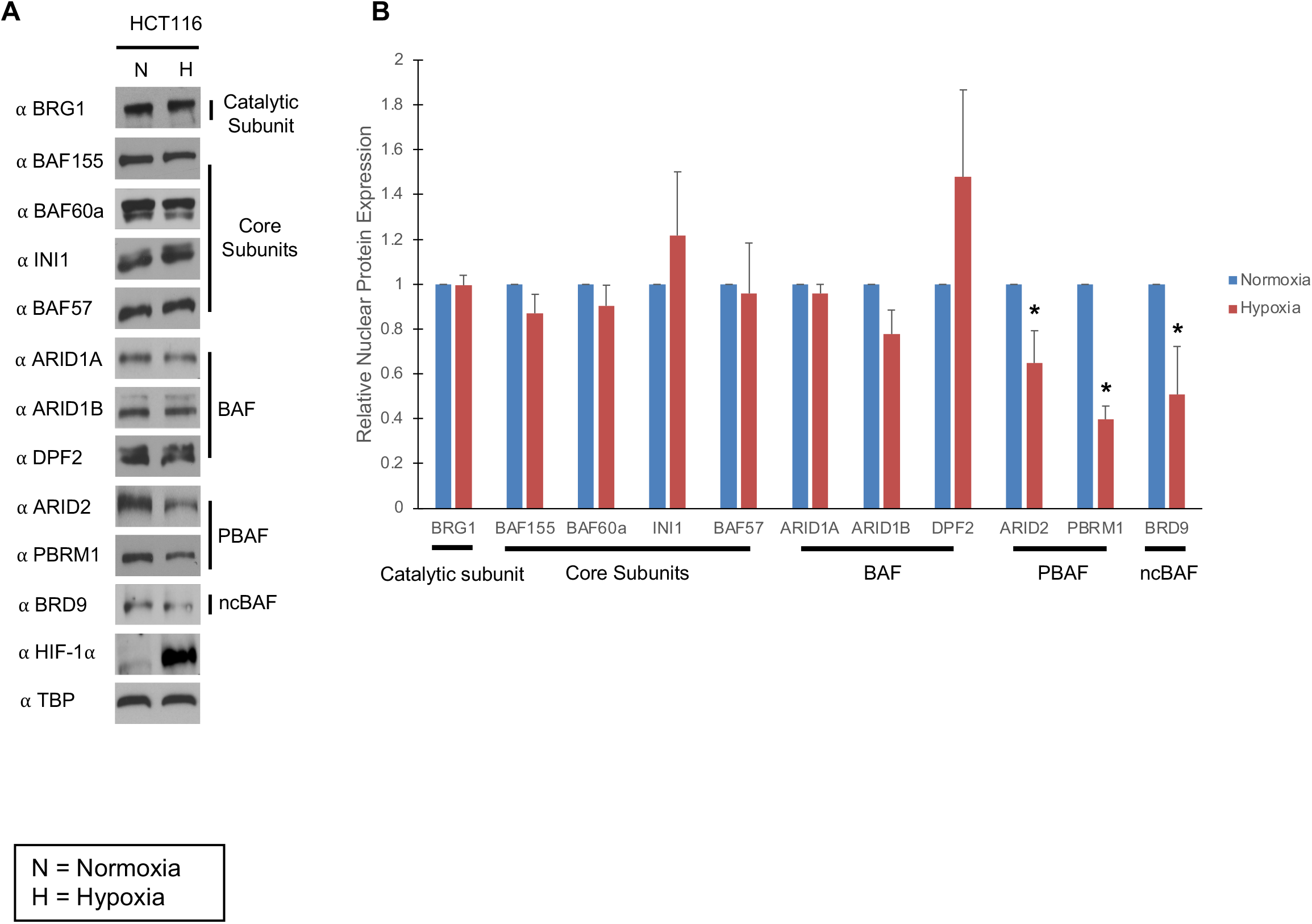
Hypoxia alters protein levels of complex specific subunits of SWI/SNF. (A) Western blot analysis of BRG1, core components (BAF155, BAF60a, INI1, and BAF57), and exclusive subunits of BAF, PBAF, and ncBAF. HIF-1α was observed to indicate hypoxic induction and TATA-binding protein (TBP) was the loading control. (B) The intensity of the bands from (A) were quantified and normalized to the corresponding loading control in normoxic and hypoxic conditions. The data plotted are averages of three independent experiments.

### Cellular composition and stoichiometry of SWI/SNF complex forms is altered during hypoxia

While our studies show that proteins levels of SWI/SNF complex specific subunits were changed in hypoxia, these changes may or may not reflect a change in stoichiometry of the entire complexes of which they are members of. To test if altered protein levels reflected changes in assembly within SWI/SNF, immunoprecipitation (IP) of endogenous SWI/SNF complexes from nuclear extracts from cells grown +/- hypoxia, were carried out with antibodies against SWI/SNF catalytic subunit BRG1, followed by mass spectrometry. We used BRG1 rather than BRM as several studies have shown that BRG1 is the catalytic subunit necessary for HIF activity and induction of several HIF1 and HIF2 dependent genes^40–44^. Also, BRG1 is present in all three SWI/SNF complexes. To assess the quality of IP, we performed a silver staining of IP’ed material (Supplementary Fig. 2A). Mass spectrometric analysis revealed that under hypoxic conditions, association of SWI/SNF core subunits including BAF155, BAF170, BAF60a, INI1, BAF57, and ACTL6A and unique subunits of BAF, ARID1A and DPF2 were retained at levels similar to those observed in normoxia (Figure 2A). However, association into BRG1 complexes by exclusive subunits of PBAF: ARID2, PBRM1, and PHF10 was significantly decreased, with moderate reduction in association of ncBAF members, BRD9 and GLTSCR1, and BAF member ARID1B (Figure 2A). Interestingly, PBAF member BRD7, did not show a significant reduction in interaction with BRG1, as observed for other PBAF specific proteins. These results were further verified by western blotting analyses of BRG1 IP samples (Figure 2B). While the decreased association of PBAF members with BRG1 was well reproduced by western blotting, the reduction in association of ncBAF members and ARID1B did not reflect similar changes observed by mass spectrometric analyses of the BRG1 IP’ed samples. These results show that hypoxia does indeed decrease association of PBAF specific members with BRG1.

**Figure 2.**
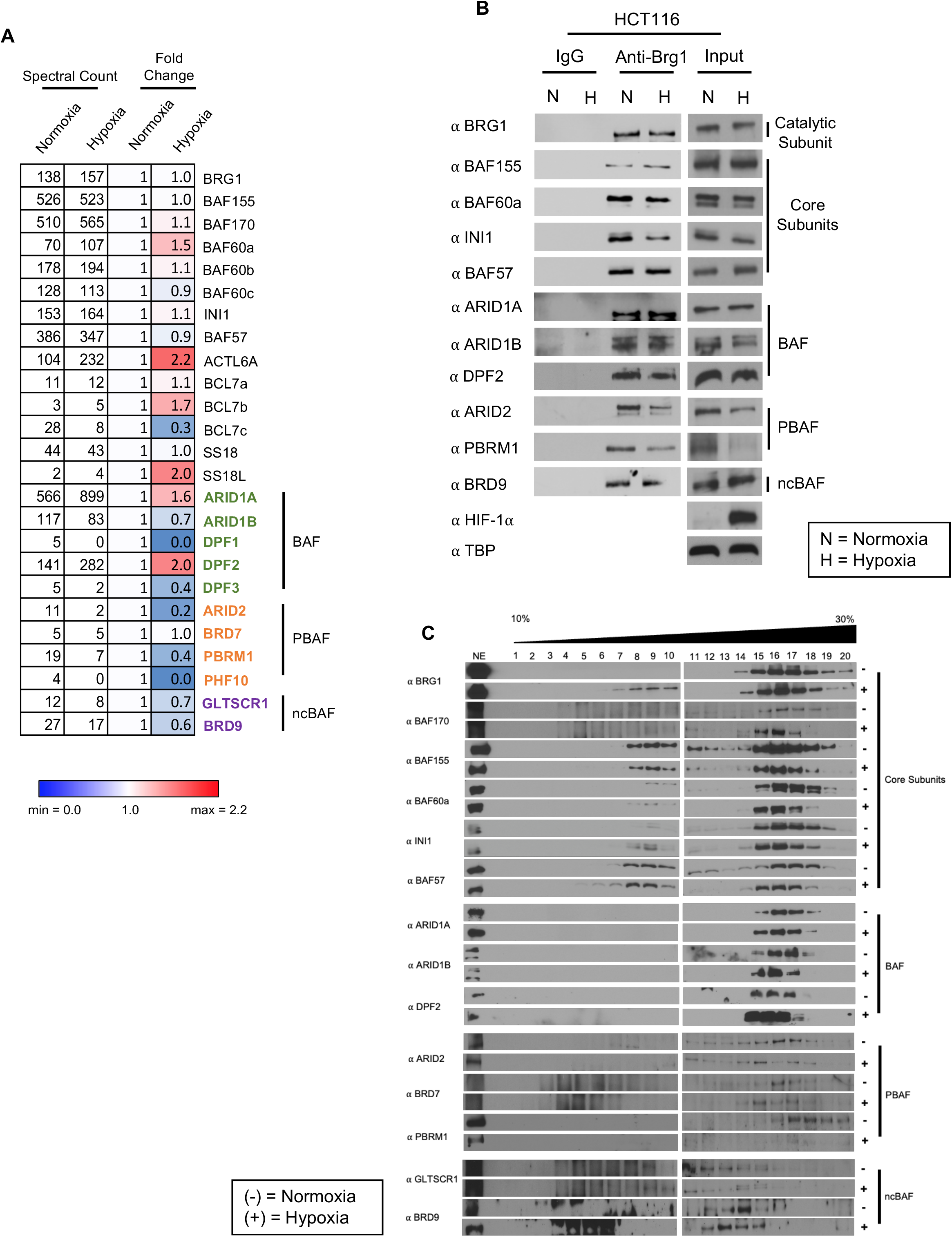
Cellular composition and stoichiometry of SWI/SNF complex forms is altered during hypoxia. (A) Immunoprecipitation-mass spectrometry of BRG1 showing SWI/SNF subunits. To compare levels of subunits, the peptide counts were first normalized to bait, BRG1, and then were normalized to their respective levels in normoxia. (B) Immunoprecipitation-western blot of BRG1 showing several core subunits and exclusive subunits of each complex. (C) Glycerol gradient of SWI/SNF complexes in normoxic (-) and hypoxic (+) conditions.

Though our results thus far provide valuable insight into each complex composition in normoxia vs hypoxia, we used a second approach of density gradient centrifugation, to delineate changes in SWI/SNF composition of individual complexes based on sizes of individual complexes. Nuclear extracts were prepared as before from cells grown +/- hypoxia, and fractionated using a 10-30% glycerol gradient. All fractions were precipitated by using trichloroacetic acid to concentrate proteins in the fractions and analyzed by western blotting with antibodies against SWI/SNF subunits. In extracts from cells grown in normoxic conditions, core components and BRG1 spread across the gradient and were associated with members of ncBAF (fractions 13-14), BAF (fractions 15-17), and PBAF (fractions 18-20). However, under hypoxic conditions, the levels of BRG1 and core components were restricted to fractions 13-14 (ncBAF) and fractions 15-17 (BAF) with a marked reduction in fractions 18-20 (PBAF) (Figure 2C). The levels of BRG1 and core SWI/SNF components aligned well with ARID1A, ARID1B, and DPF2, marking BAF in fractions 15-16 and GLTSCR1 and BRD9, marking ncBAF in fractions 13-14. However, PBAF components ARID2, PBRM1 and BRD7 observed in higher-mass fractions (fractions 18-20) in normoxic samples were reduced in hypoxic samples with PBRM1 showing the most significant reduction. With the reduction in PBRM1, the remaining PBAF subunits, ARID2 and BRD7 shifted to lower fractions, particularly fractions 15-17, suggesting a smaller complex of PBAF in the absence of PBRM1. Interestingly, we observed while loss of PBRM1 was significant, we did observe a shift in BRD7 and ARID2 to lower fractions, suggesting a smaller complex that might represent a partial PBAF complex lacking PBRM1. Additionally, we also observed BRG1 and core components of SWI/SNF in early fractions 7-10 only in hypoxia samples, which may represent partial complexes containing only the catalytic and core module of SWI/SNF, that remain after PBAF members that were lost from the complex.

As further validation, we performed IP-western blot analysis in the ES-2 ovarian carcinoma line. Western blot analysis of the IP recapitulated results observed in HCT116 cells, with a reduction in the PBAF and to a lesser degree of ncBAF member association with BRG1 and retention of BAF specific interactions (Supplementary Fig. 2B, C). Thus, while normoxic cells have all three SWI/SNF complexes, hypoxic cells retain the BAF form of SWI/SNF, and specifically reduce levels of PBAF and to a lesser degree ncBAF complexes. These results cumulatively show a fundamental alteration of stoichiometry of the various SWI/SNF complexes as a response to hypoxia and suggest that cells may require this change for gene expression and survival during hypoxic conditions.

### The BAF complex is required for induction of Hypoxic genes

It has been previously shown that SWI/SNF plays an important role in regulation of gene expression as a response to hypoxia, specifically HIF1 and HIF2 target genes^45–47^. The catalytic subunit BRG1 was required and necessary, but the requirement of individual forms of SWI/SNF complex remains unclear. Our results showing retention of BAF and to a lesser degree ncBAF forms would suggest that these complexes may play important roles in regulation of hypoxia responsive genes. To further investigate the role of the heterogeneity of the complexes in response to hypoxia in regulation of hypoxic gene expression, we knocked down unique subunits of SWI/SNF, including ARID1A, ARID1B, DPF2, PBRM1, BRD9 and control (non-target), by small interfering RNA (siRNA). Knock down of subunits was verified by western blotting (Supplemental Fig. 3A). We first examined transcript levels of ANGPTL4 and CA9, HIF1 target genes that have been shown to be dependent to SWI/SNF for their expression^43^. We found that knockdown of ARID1A and DPF2 markedly reduced the hypoxic induction of ANGPTL4, with no significant changes in ANGPTL4 levels on loss of ARID1B, PBRM1 or BRD9 (Figure 3A). In the case of CA9 while loss of ARID1B reduced its induction, loss of ARID1A and DPF2 did not produce significant affects (Figure 3B). Interestingly, loss of PBRM1 and BRD9 increased expression, suggesting repressive roles of PBAF and ncBAF. Next, we examined levels of PAI1, a HIF2 target gene that is also known to be SWI/SNF dependent^43^. We found that knockdown of all BAF specific subunits reduced expression of PAI1, but loss of BRD9 increased expression of this gene (Figure 3C). ARID1A and ARIB1B are assembled into distinct BAF complexes and are not found simultaneously with a single BAF complex. These results suggest that while some hypoxic genes can be regulated by a unique form of BAF, the BAF complex forms may have overlapping roles in expression of others. Further, the increase in levels of mRNA of hypoxia responsive genes on loss of BRD9, suggest a suppressive role of ncBAF in hypoxic gene expression. Our results mirror studies which have shown that loss of BRD9 in fact increases levels of transcripts of hypoxia responsive genes^33^.

**Figure 3.**
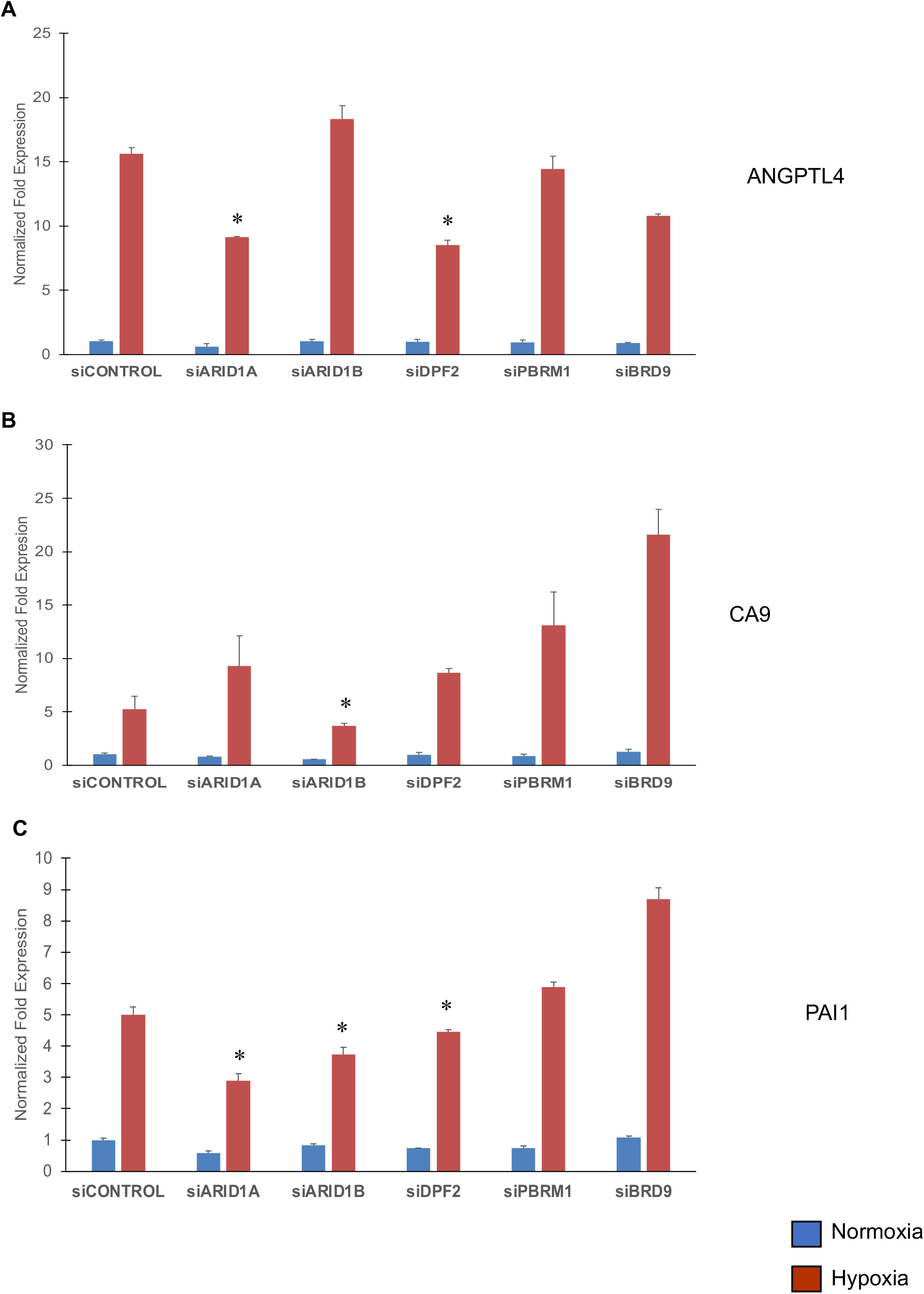
The BAF complex is required for induction of Hypoxic genes. HCT116 cells were transfected with ARID1A, ARID1B, DPF2, PBRM1, BRD9 and Control siRNA. (A) qRT-PCR analysis of ANGPTL4. (B) qRT-PCR analysis of CA9. (C) qRT-PCR analysis of PAI1. The data plotted are averages of three independent experiments. The data was normalized to non-target control normoxia sample, that was set to 1.

### BAF complex members ARID1A and DPF2 are important for survival during hypoxia

The retention of BAF complex during hypoxia and its role in expression of hypoxia responsive genes, suggests a possible role of this complex in cell survival during hypoxia. To further explore the importance of the BAF and other forms of SWI/SNF in cell survival during hypoxia, we examined viability of cells lacking ARID1A, ARID1B, DPF2, PBRM1, and BRG1 by performing an MTT assay. Subunits were targeted by siRNA as described above and non-target siRNA was used as control (Supplemental Fig. 3A). We observed that depletion of the ATPase subunit, BRG1, significantly reduced cell viability both in normoxia, as well as, in hypoxia (Figure 4). Loss of BRG1 has been shown previously by others to be required for cell proliferation^47–50^, thus loss of viability is expected, as loss of a catalytic subunit will significantly affect SWI/SNF ability to alter chromatin structure needed for altered gene expression. Knockdown of ARID1A, ARID1B, PBRM1 and BRD9 under normoxic conditions showed no significant reduction in cell viability but interestingly, we observed an increase in cell proliferation when ARID1B and PBRM1 were depleted (Figure 4). However, although loss of ARID1A under normoxic conditions showed no changes in cell viability as compared to the normoxic control, we observed that cells were more sensitive to ARID1A loss in hypoxic conditions (Figure 4A). We also observed knockdown of DPF2 decreased cell viability in hypoxic conditions, however ARID1B was not necessary for cell survival (Figure 4A). Loss of PBRM1 and BRD9 did not show significant effects on hypoxic survival. Our results point to the fact that ARID1A/DPF2 within BAF play critical roles in response to hypoxic stress. While loss of ARID1B would be expected to show some effect on cell survival given its role in hypoxic gene expression, the roles of ARID1B could be compensated by ARID1A, which is more critical for hypoxic response. In conclusion, an ARID1A/DPF2 containing BAF complex is necessary under hypoxic conditions and that loss of this complex affects gene expression and cell survival.

**Figure 4.**
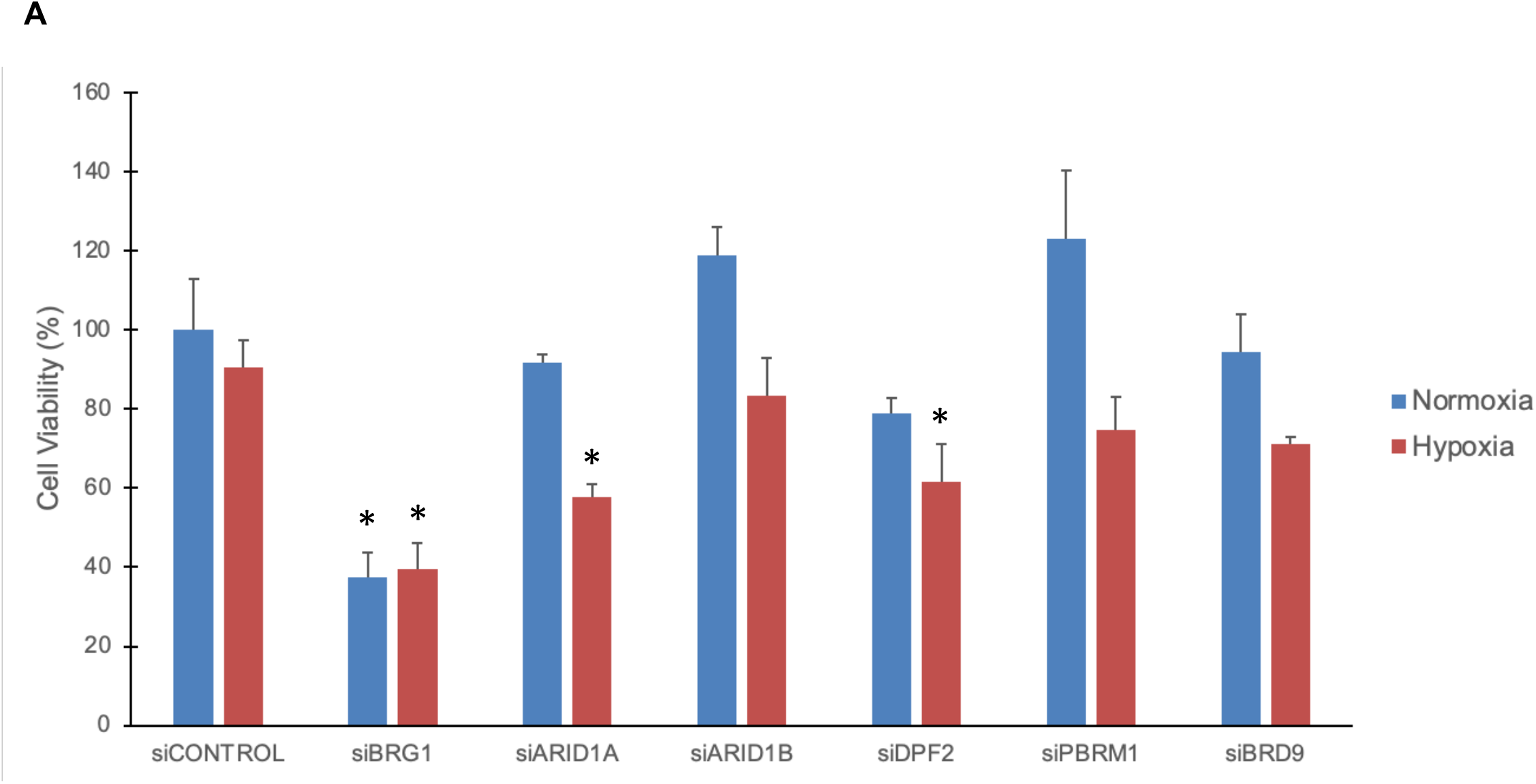
BAF complex members ARID1a and DPF2 are important for survival during hypoxia. (A) HCT116 cells were more sensitive to ARID1A and DPF2 loss in siRNA knockdown. Cells were grown in normoxia or hypoxia, following KD of subunits and cell viability was measured using MTT assay. The data plotted are averages of three independent experiments. The data was normalized to non-target control normoxia sample, that was set to 100%.

## Discussion

There are three well defined forms of the SWI/SNF complex, BAF, PBAF and ncBAF. These complexes share some common subunits, including BRG1, the catalytic subunit, and core subunits, BAF155 and BAF60a, but the identity and genomic functions of each complex is represented by their unique subunits. In this study, we have evaluated how the cells make use of the different forms of SWI/SNF in order to respond to hypoxia.

The role of SWI/SNF in hypoxia has been demonstrated through the catalytic subunits, BRG1 and BRM. Studies have shown that BRG1 and/or BRM regulate gene expression of several HIF-1α and HIF-2α genes and loss of these subunits, significantly reduces induction of hypoxia responsive genes ^40,43^. BRG1 is also necessary for cell proliferation and migration during hypoxic response^51^. As mentioned previously, BRG1 is found in all three complexes, and the common theme shows SWI/SNF to be critical in hypoxic response by BRG1, but the identity of the specific form/forms of SWI/SNF that plays critical roles in hypoxia mediated gene expression and cell survival remains unknown. Our studies show that as a response to hypoxia, cells regulate the abundance of various forms of SWI/SNF, thereby altering the balance of the complexes as summarized in Figure 5. In normoxic conditions, we observed all three complexes were present but in hypoxic conditions, there was a shift to an ARID1A/DPF2-containing BAF complex while PBAF and ncBAF were downregulated, as observed by reduction in levels of ARID2 and PBRM1, and BRD9, and their reduced association with BRG1. Interestingly, we also observed a possibly smaller complex of PBAF from the glycerol sedimentation assay (Figure 2C). PHF10 peptide counts were non-existent from mass spectrometry and studies have reported the instability of the subunit^52^. The reduction in protein levels of PBRM1 and PHF10 (Figure 1A/B), and their decreased association with PBAF complex (Figure 2B, 2C), suggests that cells may need their loss for hypoxic response. The mechanism of reduction of PBRM1 and consequence of retention of PBRM1 in hypoxic cells warrants further scrutiny.

**Figure 5.**
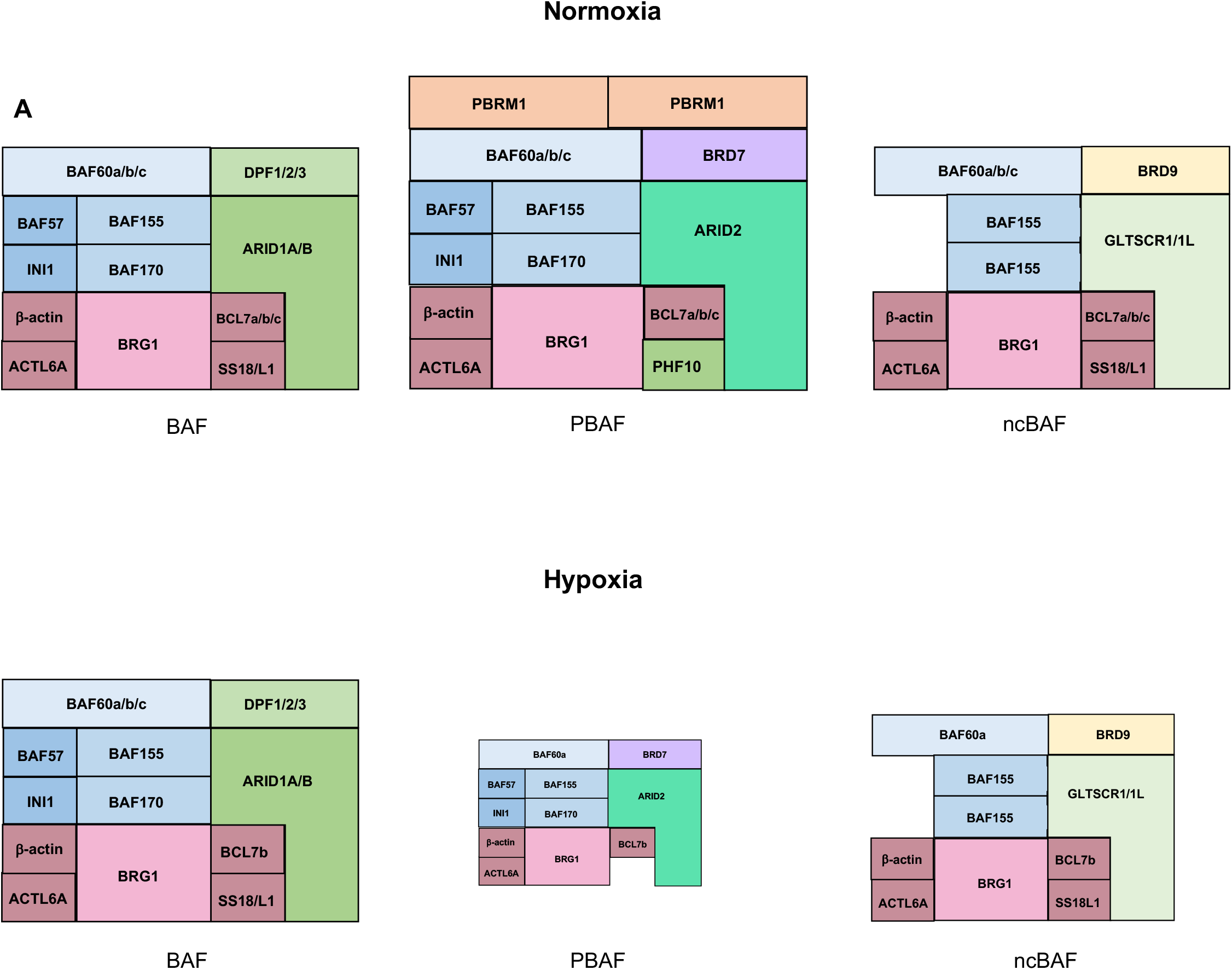
Model for how the heterogeneity of SWI/SNF shifts from normoxia to hypoxia. Under normoxic conditions, all three complexes are present. However, when the condition becomes hypoxic, where the oxygen level drops from 21% to 1% O_2_, we observed an ARID1A/DPF2 containing BAF complex played an important role. The PBAF complex lost PBRM1 and PHF10 subunits, becoming a smaller complex and ncBAF was reduced in terms of its presence.

Our studies further show that retention of BAF complex during hypoxia, is needed for induction of hypoxia responsive genes, including ANGPTL4, CA9 and PAI, induction of which is reduced when cellular levels of ARID1A, ARID1B, or DPF2 are reduced. An interesting observation is that knock down of BRD9 showed increased induction of CA9 and PAI1, indicating BRD9 may play a suppressive role in induction of hypoxic genes. We showed that although ncBAF is downregulated, the complex is not completely loss and suggests that the complex maintains a presence, most likely to modulate levels of hypoxic mediated gene expression and perhaps suppression of other genes. We also observed that knockdown of ARID1A or DPF2 significantly decreased cell viability on hypoxic stress. These findings provide evidence that ARID1A and DPF2 containing BAF plays a significantly role not only for gene expression but survival during hypoxic conditions.

In summary, our study provides evidence that in normoxic conditions, all three SWI/SNF assemblies are present but when the environment becomes hypoxic, ARID1A/DPF2-containing BAF complex is important for hypoxia mediated gene expression of several HIF1 and HIF2 genes and for cell viability during hypoxic stress. Furthermore, we also report a possible smaller PBAF complex that does not contain PHF10 and PBRM1. Therefore, our data provides evidence that cells dynamically alter the heterogeneity of the complexes in order to respond to changes in cellular environments.

## Materials and Methods

### Cell lines

HCT116 cell line was obtained from Horizon Discovery. ES-2 cell line was obtained from ATCC. HCT116 and ES-2 were maintained in McCoy’s Medium (Gibco) containing 10% FBS (Gibco), supplemented with 1% penicillin and streptomycin (Gibco). Cell lines were routinely tested for mycoplasma. Cells were cultivated at 37°C in 5% CO_2_, in accordance with the supplier’s instructions, unless otherwise noted.

### Hypoxic Cell Culturing

The StemCell Technologies Hypoxia Incubator Chamber was used in this study for the generation of a hypoxic environment. After 24-hour cultivation in conventional cell culture incubator (allowing cells to attach to plates), plates were transferred to the chamber and 1% O_2_, 5% CO_2_, and 94% N_2_ gas mixture was purged into the chamber and placed into the conventional cell culture incubator for 24-hour incubation, unless otherwise noted.

### Nuclear extract preparation

Cells were seeded into 15 cm^2^ dishes with 8 × 10^6^ cells. After 24 hours, cells were incubated in normoxic or hypoxic conditions as mentioned previously. Cells were harvested in ice-cold PBS. Cytoplasmic extract was prepared in Buffer A (10 mM HEPES, pH 7.9, 1.5 mM MgCl_2_, 10 mM KCl, 0.1 mM EDTA) with complete protease inhibitor and 1 mM PMSF. Samples were kept on ice for approximately 30-45 minutes with gentle mixing in between rupture cells. Samples were spun at 5000 g for 5 minutes at 4°C to pellet nuclei. Supernatant contained cytoplasmic extract. Nuclear extract was prepared in Buffet B (20 mM HEPES, pH 7.9, 1.5 mM MgCl_2_, 420 mM KCl, 0.2 mM EDTA, 25% glycerol, 1% Triton X-100) with complete protease inhibitors, 0.1 mM DTT, 1 mM PMSF, benzonase and heparin. Extract was left at room temperature for 15 minutes to rotate end over end. Extract was moved to 4°C for 1 hour to rotate end over end. Extract was spun at 15,000 rpm (benchtop centrifuge) for 30 minutes at 4°C. Protein concentration was determined by BCA (Thermo Fisher). Extracts were subjected to western blot analysis, immunoprecipitation, or glycerol sedimentation assay.

### Immunoprecipitation

Approximately 2.5 µg of BRG1 antibody (Abcam) and 20 µl of Protein A Dynabeads (Invitrogen) were crosslinked at 4°C, rotating end over end for 4 hours. Antibody-bound beads were washed 4X with ice cold PBS and once with 1 ml 0.1 M sodium borate (pH 9.0). Antibody-bound beads were resuspended with sodium borate with the addition of 5.2 mg of dimethylpyrimilidate (DMP) and allowed to rotate end over end at room temperature for 30 minutes. Antibody-bound beads were washed once with 0.2 M ethanolamine (pH 8.0) and then once more and allowed to rotate end over end at 4°C overnight, prior to nuclear extraction. Nuclear extract preparation was used for IP. Approximately 1 mg of protein extract was added to Buffer C (20 mM HEPES, pH 7.9,1.5 mM MgCl_2_, 300 mM KCl, 0.2 mM EDTA, 10% glycerol, 0.1% NP-40) with complete protease inhibitors, 0.1 mM DTT and 1 mM PMSF and incubated on ice for 15 minutes and spun at 14,000 rpm for 20 minutes at 4°C. Supernatant was removed and added to antibody-bound beads and incubated at 4°C, rotating end over end, for 4 hours. Nuclear extracts were eluted in 0.1 M glycine (pH 2.5) for 2 minutes on ice with constant flicking of tube. Tris-HCl pH 8.8 was added to neutralize the pH.

To verify quality of samples, silver staining and western blotting was performed. The SDS-PAGE was fixed in Buffer 1 (50% ethanol, 10% acetic acid and 40% deionized water) overnight and washed 3X for 15 minutes each thereafter. Gels were sensitized in Buffer 2 (0.02 g of Na_2_S_2_O_3_ in 100 ml of deionized water) for 25 minutes, washed 3X with deionized water and allowed to develop in Buffer 4 (6 g of Na_2_CO_3_, 2 ml of Na_2_S_2_O_3_, 50 µl of formaldehyde in 100 ml of deionized water) until a desired signal and stopped by stop solution (10% acetic acid in 100 ml of deionized water) for 20 minutes.

### Mass Spectrometry

Endogenous complexes were purified from IP protocol. Complexes were TCA-precipitated and then urea-denatured, reduced, alkylated, and digested with endoproteinase Lys-C (Roche) followed by modified trypsin (Promega). IP samples was sent to Stowers Institute for Medical Sciences, Proteomics Center.

### Glycerol sedimentation gradient

Nuclear extracts were placed on top of a 10 ml 10-30% glycerol gradients containing 25 mM HEPES, pH 7.9, 0.1 mM EDTA, 12.5 mM MgCl_2_, 100 mM KCl and supplemented with complete protease inhibitors, 1 mM DTT and 1 mM PMSF. Tubes were placed into SW41 rotor and centrifuged at 40,000 rpm for 16 hours at 4°C. 500 µl fractions were collected from the top of the gradient. 500 µl of 100 mM Tris-HCl, pH 8.5 and 250 µl TCA were added to fractions, inverted, and placed into 4°C, overnight. Factions were spun at 14,000 rpm for 30 minutes at 4°C, supernatant removed, leaving approximately 20 µl and washed twice with 500 µl of cold acetone, inverting gently. Supernatant was removed after washes and allowed to air dry in the hood. Fractions were then used for western blot analysis.

### Western blot analysis

For western blot analysis, samples were separated on a 10% SDS-polyacrylamide gel electrophoresis gel and transferred onto PVDF membrane. Membranes were blocked in 5% milk with TBST and incubated with primary antibodies overnight. Membranes were washed 3X with TBST and incubated with secondary antibodies for 1 hour. The following primary antibodies used were: anti-BRG1 (Abcam, catalog#: ab110641), anti-BAF170 (Bethyl, catalog#: A301-039A), anti-BAF155 (Bethyl, catalog#: A301-020A), anti-BAF60a (Santa Cruz, catalog#: sc-514400), anti-INI1 (Abcam, catalog#: ab192864), anti-BAF57 (Bethyl, catalog#: A300-810A), anti-ARID1A (Sigma, catalog#: HPA005456), anti-ARID1B (Bethyl, catalog#: A301-046A), anti-ARID2 (Bethyl, catalog#: A302-229A), anti-BRD7 (Proteintech, catalog#: 51009-2-AP), anti-PBRM1 (Sigma, catalog#: HPA015629), anti-GLTSCR1 (GenScript), anti-BRD9 (Bethyl, catalog#: A303-781A) anti-HIF-1α (Bethyl, catalog#: A300-286A), and anti-TBP (Proteintech, catalog#: 66166-1-Ig).

### siRNA Transfections

300,000 cells were seeded into 6 well plates and incubated overnight. A double transfection was performed using an siRNA cocktail (ON-TARGET Plus Pool, Horizon Discovery). 10 µl of 5 µM siRNA was diluted into 250 µl Opti-MEM (Gibco) and 10 µl of Lipofectamine 3000 (Invitrogen) was diluted into 250 µl of Opti-MEM. Solutions were incubated at room temperature for 5 minutes and then lipofectamine mix was added to siRNA mix and incubated at room temperature for 20 minutes. The mixture was added dropwise in each well and incubated at 37°C overnight. Another round of transfection was performed 24 hours later. The cells were harvested and seeded for protein analysis, RNA isolation, and MTT assay.

### MTT Assay

5,000 transfected siRNA cells were seeded into 200 µl of media and cultured in 96 well plates. Cells were incubated in either normoxic or hypoxic conditions for 24 hours. After, 5 mg/ml of MTT (Sigma) was added and incubated at 37°C for 4 hours before 200 µl of acidified isoproponal was added to each well and placed on a shaker for 20 minutes, covered in foil and absorbance was determined using a spectrophotometer at a wavelength of 490 nm (BioTek).

### RNA isolation and reverse transcriptase-quantitative PCR analysis

Total RNA was extracted from cells using Trizol (Invitrogen). 1-2 µg of RNA was reversed transcribed into cDNA using SuperScript IV First-Strand cDNA Synthesis Reaction according to manufacturer’s instructions. qRT-PCR was performed using the Roche Light Cycler 480 II and SYBR Green (Applied Biosystems). The data used housekeeping 18S rRNA and β-actin gene for normalization. The expression of CA9, ANGPTL4 and PAI1 was analyzed using specific primers (Table 1). The expression levels of the above genes were calculated as ΔΔCT, where CT is cycle threshold, normalizing to normoxic control samples.

**TABLE 1.**
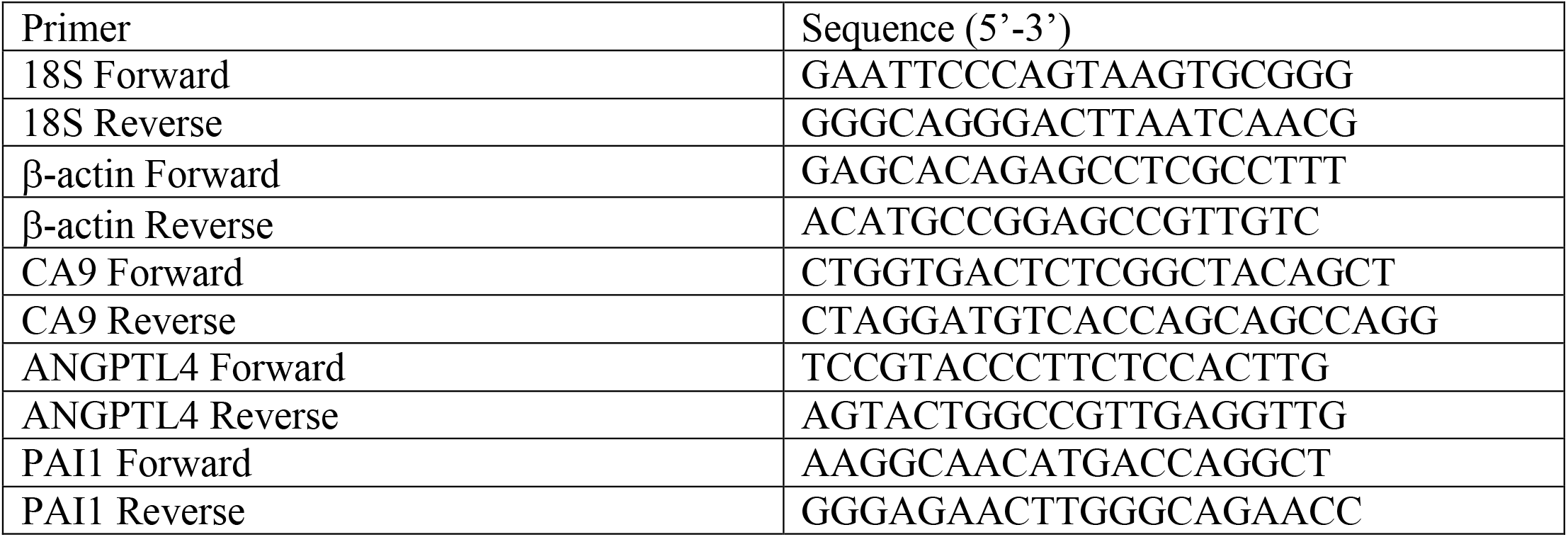
Primer sequences used for qRT-PCR

## Supporting information

all supplemental figures

## Acknowledgements

We would like to thank all members of the Dutta lab for their help and comments. This work is supported by funding to AD from the Rhode Island Foundation Medical Research Grant, NIH/NIGMS P20GM103430/Rhode Island IDeA Network for Biomedical Research Excellence (RI-INBRE) and the University of Rhode Island.

---

Supplemental Figure 1 – (A) Western blot analysis in ES-2 cells +/- hypoxia for BRG1, core components (BAF155, BAF60a, INI1, and BAf57), and exclusive subunits of BAF, PBAF, and ncBAF. HIF-1α was observed to indicate hypoxic induction and TATA-binding protein (TBP) was the loading control. (B) The intensity of the bands from Supplemental Figure 1A were quantified and normalized to the corresponding loading control in normoxic and hypoxic conditions. The data plotted are averages of three independent experiments.

Supplemental Figure 2 –Silver staining of BRG1 IP from HCT116 cells (A) and ES2 cells (B). (C) Immunoprecipitation-western blotting analysis of BRG1-IP for several core subunits and exclusive subunits of each complex in ES-2 cells grown +/- hypoxia.

Supplemental Figure 3 – (A) Western blot analysis of siRNA knockdown of ARID1A, ARID1B, PBRM1, BRD9 and Control-non target in HCT116 cells.

