## Supplementary material for "Hypoxic response is driven by the BAF form of SWI/SNF": all supplemental figures

### Supplemental information

**Supplemental Figure 1**

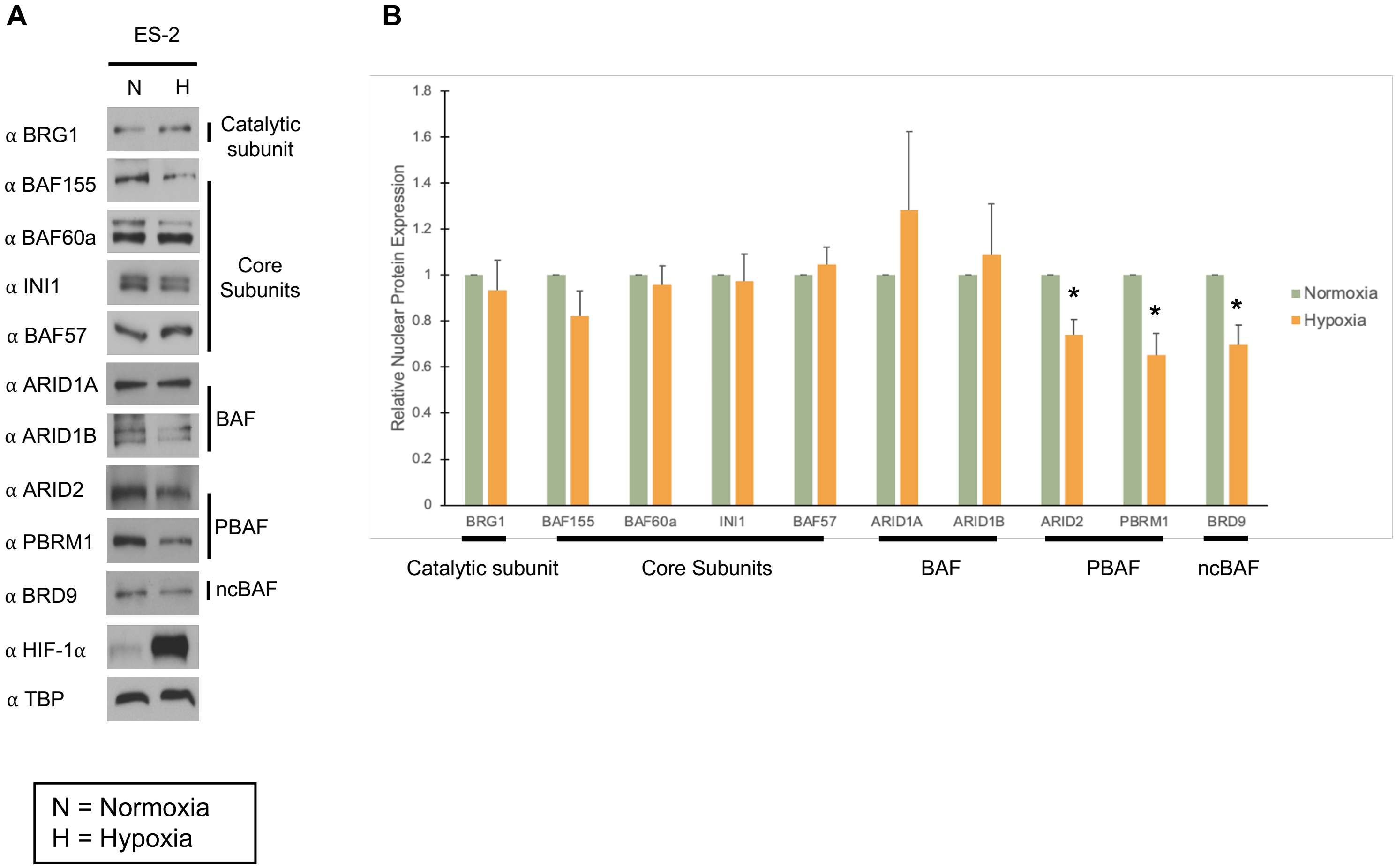


Supplemental Figure 1 – (A) Western blot analysis in ES-2 cells +/- hypoxia for BRG1, core components (BAF155, BAF60a, INI1, and BAf57), and exclusive subunits of BAF, PBAF, and ncBAF. HIF-1α was observed to indicate hypoxic induction and TATA-binding protein (TBP) was the loading control. (B) The intensity of the bands from Supplemental Figure 1A were quantified and normalized to the corresponding loading control in normoxic and hypoxic conditions. The data plotted are averages of three independent experiments.

**Supplemental Figure 2**

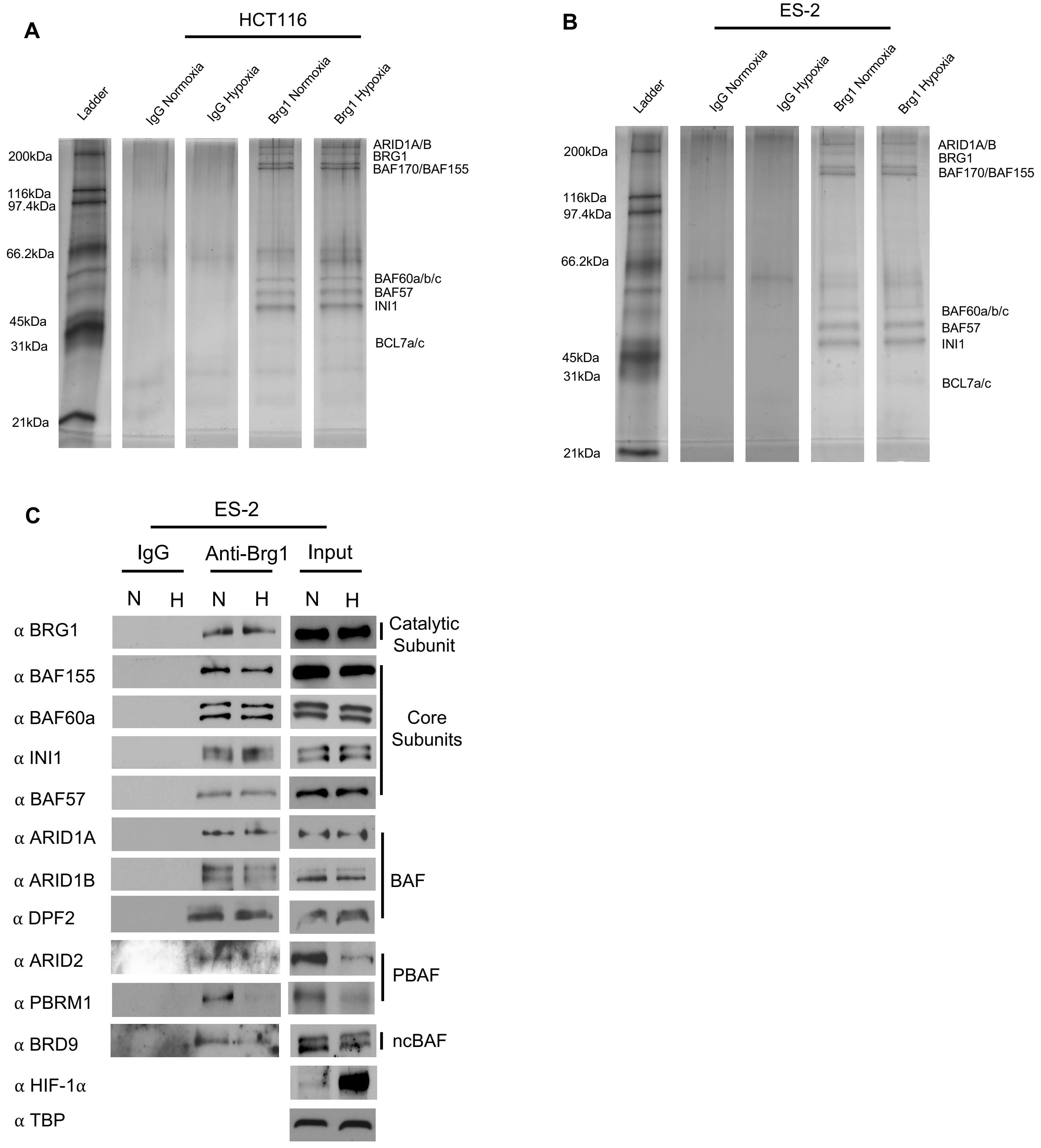


Supplemental Figure 2 –Silver staining of BRG1 IP from HCT116 cells (A) and ES2 cells (B). (C) Immunoprecipitation-western blotting analysis of BRG1-IP for several core subunits and exclusive subunits of each complex in ES-2 cells grown +/- hypoxia.

**Supplemental Figure 3**

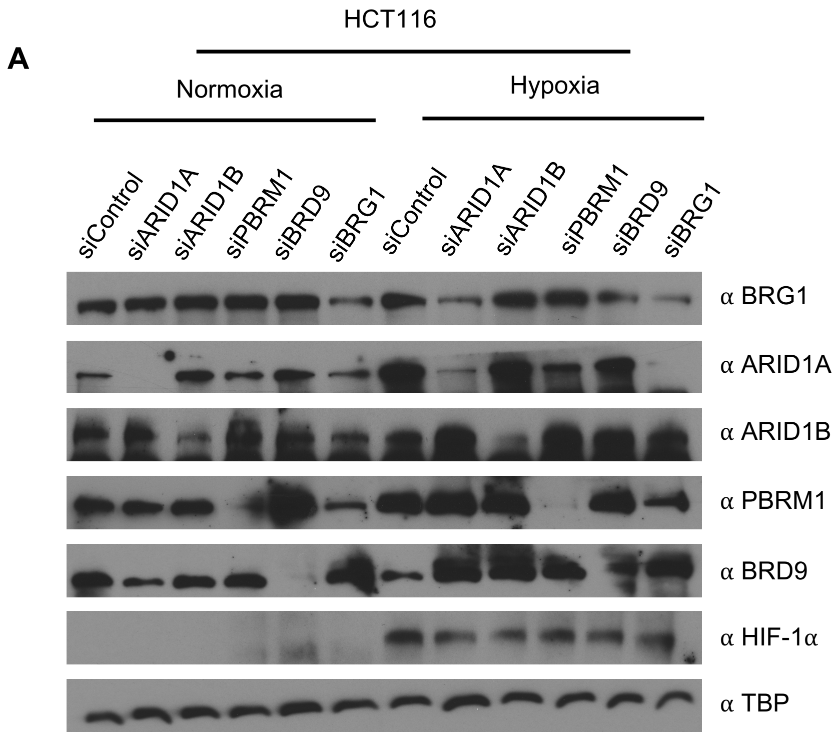


Supplemental Figure 3 – (A) Western blot analysis of siRNA knockdown of ARID1A, ARID1B, PBRM1, BRD9 and Control- non target in HCT116 cells. TATA-binding protein (TBP) is used as loading control.
